# DNA virus–host patterns in lake and marine environments over the last glacial cycle

**DOI:** 10.1101/2025.06.18.660323

**Authors:** Christiane Boeckel, Simeon Lisovski, Kathleen R. Stoof-Leichsenring, Josefine Friederike Weiß, Sisi Liu, Lars Harms, Ulrike Herzschuh

## Abstract

Viruses are crucial to population control, biogeochemical cycling, and host evolution, making them essential for ecosystem function. Here, we explore long-term virus–host interactions in lake and marine environments across the late Pleistocene and Holocene.

We analysed sedimentary ancient DNA (sedaDNA) from five Siberian lakes and four Subarctic/Antarctic marine cores to infer past DNA virus taxa from metagenomic sequences. Viruses accounted for 0.089% (357 161 reads) of mapped reads across 2084 taxa. Virus communities differ between lakes and marine sites, with lakes dominated by *Caudoviricetes* and marine environments featuring *Caudoviricetes* and *Algavirales*. Each time series shows compositional changes in virus communities from the Pleistocene to the Holocene, supporting the inference that sedaDNA can reconstruct time-resolved ancient viral assemblages.

Among the most abundant viruses, we identified 83 virus–host pairs known from published literature to infect bacterial, archaeal, or eukaryotic hosts, and assessed their associations using co-variation patterns. Over millennia, virus–host co-variations are particularly stable in marine systems, especially for phytoplankton-infecting viruses. However, in the Bering Sea, we find a lack of virus–host correlation, likely because an Arctic Pelagibacter strain expanded rapidly after the Bering Strait opened, potentially due to absent viral infection.

Antagonistic patterns also appear between bacteriophages and hosts, possibly linked to shifts between lytic and lysogenic cycles in response to environmental changes.

This study shows that sedaDNA can reveal ancient viral community structures and long-term ecological patterns, highlighting the value of ancient viromes in understanding ecosystem-specific responses to environmental change.

## Introduction

Viruses are critical components of ecosystems but are often overlooked in biodiversity assessments. Their reliance on hosts for replication affects host fitness and can cause entire community shifts. Beyond controlling host populations, viruses shape host evolution via gene transfer and are involved in biogeochemical cycling [1–3]. Understanding long-term virus– host dynamics is therefore crucial to predicting how ecosystems respond to environmental change.

Viruses infecting bacteria (bacteriophages) use host cellular machinery to reproduce, ultimately causing cell lysis. However, some bacteriophages enter a lysogenic life cycle, integrating into the host genome as proviruses and remaining dormant until conditions favour viral production and a switch to lysis [4,5]. Similar latent phases occur in some eukaryotic viruses [6,7]. This dormancy allows viruses to form "seed banks" within ecosystems, re-emerging when conditions favour replication [8]. Periods of reduced viral production may also result from the evolutionary arms race between viruses and hosts, described by the Red Queen hypothesis. In this constant struggle, hosts evolve mechanisms to resist infection, while viruses adapt to overcome these defenses, leading to suppressed viral replication while hosts gain a temporary advantage [9]. The mechanism of how potential host resistance or viral dormancy affects virus–host dynamics resulting in reduced virus productivity over long timescales is poorly understood.

Viruses show strong habitat specificity, with distinct communities found in different environments [10]. Yet within a given habitat, viral populations can be widely distributed, as seen in marine viruses dispersed by ocean currents [8,10]. In contrast, freshwater virus communities, such as those in polar lakes, are more isolated and distinct [11]. Although these patterns are documented in modern systems, it remains unclear whether such ecological structures persist over geological timescales or how they respond to millennial scale, natural environmental changes.

Temperature is a major driver of viral community composition and infection dynamics in both marine [8,12] and freshwater systems [11], which makes them susceptible to global warming [13,14]. The Late Quaternary was characterised by significant temperature variations of several degrees Celsius [15]. During the last glacial period, the Northern Hemisphere was extensively glaciated, reaching its maximum during the last glacial maximum (LGM; 26.5–19 ka BP) [15]. Yet, despite the extremely cold climate, Northeast Siberia remained mostly ice-free [16]. Glaciation caused sea levels to drop by 120–130 metres [15], resulting in the exposure of a land bridge between Siberia and Alaska, known as Beringia, which disrupted water flow between the Arctic Ocean and the Northern Pacific. As global temperatures began to rise around 19 ka BP [15], progressive deglaciation and sea-level rise gradually submerged the land bridge, leading to the reopening of the Bering Strait around 11-12 cal ka BP [17–19]. A particularly intense period of warming began around 14 ka BP in the Northern Hemisphere, marking the transition from the last glacial period of the late Pleistocene towards the Holocene interglacial [20]. Meanwhile, the Southern Hemisphere experienced a temporary cooling between 14.7 and 12.7 ka BP, known as the Antarctic Cold Reversal (ACR) [21]. During this period, atmospheric and oceanic temperatures dropped, expanding sea ice around the Antarctic continent [22]. Such major environmental changes in both the Northern and Southern Hemisphere likely shaped virus–host dynamics, but direct evidence of how viral communities responded is lacking.

Past virus–host interactions have to be inferred from environmental records. Well-preserved remains from vertebrates enable the investigation of ancient host-associated viruses.

Ancient viruses have been identified in preserved feces [23], bones [24], and soft tissues [25,26]. These samples offer insights into viral interactions with animals, humans, and their respective microbiomes at the time of their decease. Ancient environmental biomes and their viral assemblages at the time have been inferred from ancient DNA in permafrost soil [27–30] and glacier ice [31,32], some of which covered transitions between glacial and interglacial periods [27,32]. However, reconstructions of ancient viral communities in aquatic systems and time-resolved records are missing so far.

In the last two decades, sedimentary ancient DNA (sedaDNA) has emerged as a reliable proxy to reconstruct ecosystem composition of the past across all domains [33,34]. DNA from local and catchment aquatic and terrestrial organisms is buried and preserved in the sediments. PCR-independent metagenomic shotgun sequencing of the sedaDNA provides a comprehensive picture of the source organisms and thus enables the simultaneous investigation of the viral community and its hosts within the same dataset [27]. This method is especially useful for viruses, which lack a universal marker gene and cannot be studied with metabarcoding [35]. However, whether sedaDNA can be reliably used to reconstruct ancient viral communities and reveal long-term virus–host dynamics remains untested.

Here, we investigate the DNA virus community preserved in sediments from marine and lake environments across the Late Quaternary. Using metagenomic data from eight sediment cores spanning Subarctic, Arctic, and Antarctic sites, we aim to elucidate the spatial and temporal patterns of ancient viral communities. Specifically, we (1) compare viral assemblages between lake and marine systems and across the last glacial and current interglacial, and (2) examine virus–host co-occurrence patterns to investigate ecological interactions and potential perturbations in the past. In this study, we demonstrate that sedaDNA enables the reconstruction of ancient viral communities and virus–host dynamics in both lake and marine environments over millennial timescales. We identified distinct differences in viral community composition between lakes and marine systems, with environmental context (lake vs. marine) exerting a stronger influence than geological epoch. Temporal patterns in viral taxa reflect major climatic transitions, particularly the shift from the last glacial period to the Holocene. Additionally, we find both positive and negative long-term correlations between viruses and their hosts, suggesting that periods of reduced viral production, potentially driven by environmental change, host resistance, or viral dormancy, shaped virus–host interactions over time.

## Materials and methods

### Sediment cores and DNA extraction

In this meta-analysis, five Siberian lake sediment cores (Levinson-Lessing, Lama, Ilirney, Ulu, Bolshoe Toko) were compared to three marine cores from the Northern Pacific (SO201- 2-KL77, SO201-2-KL12) and the Bransfield Strait, Antarctic Peninsula (PS97/072-01). These cores generally span the Pleistocene-Holocene, with the oldest record dating back to 124 cal ka BP. Their collection, dating and age-depth models are described in previous publications (Table 1). The sedaDNA was extracted using the DNeasy PowerMax Soil Kit (Qiagen, Germany) with a modified protocol described in detail in [36]. In total, 418 samples and 133 extraction blanks were produced. After DNA extraction, samples and blank controls were concentrated and purified using the GeneJet PCR Kit (ThermoFisher) and 10 µL of a 3 ng/µL solution of concentrated sample sedaDNA (and 10 µL DNA from the extraction blanks) were used as input to the single-stranded library preparation protocol [37,38]. Each library batch contained, on average, six sedaDNA samples, one extraction blank, and one library blank.

**Table 1:**
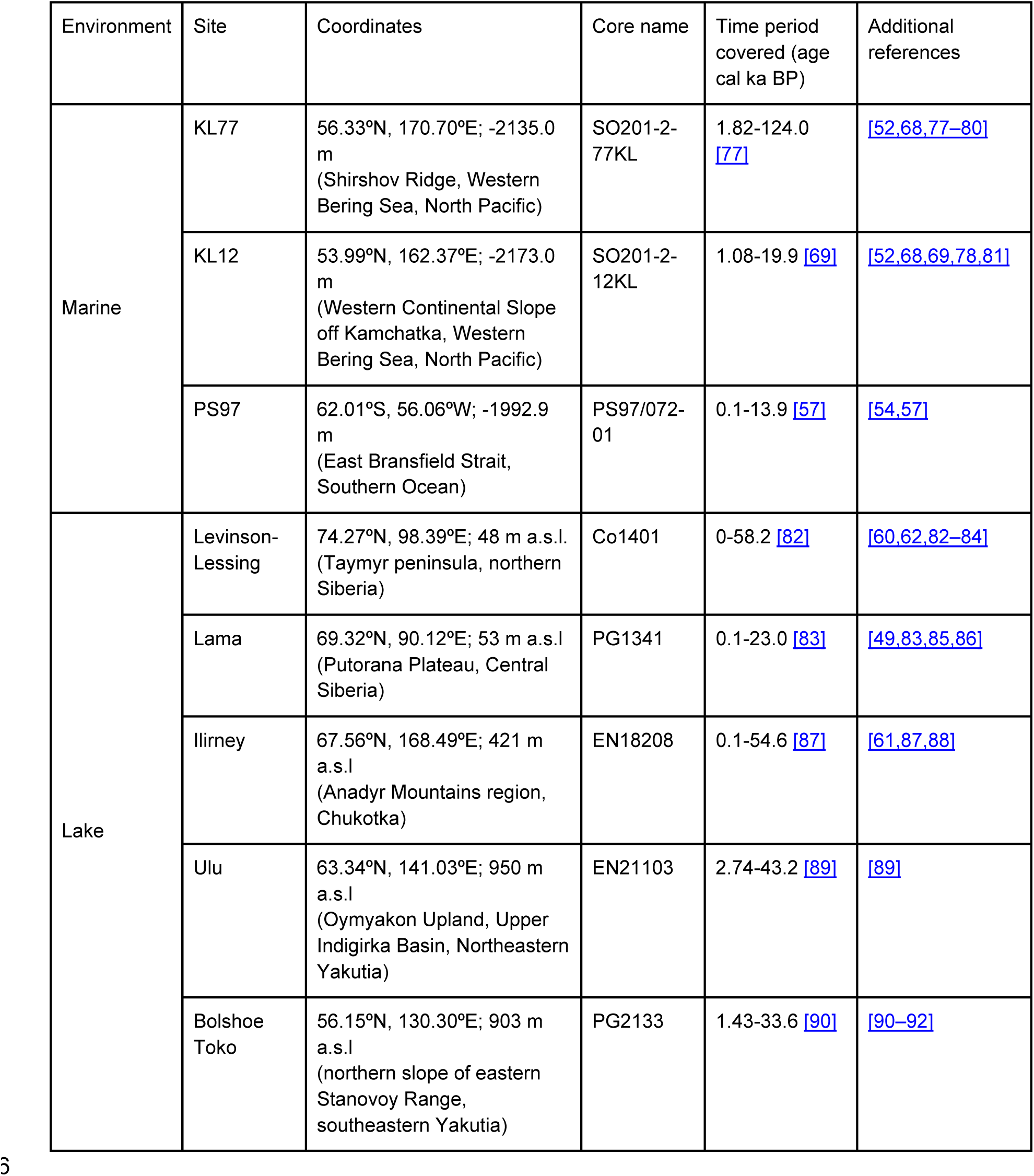
Metadata of the sampled sites.

The full description of the library preparation and shotgun sequencing is described in [34]. Further details about the exact numbers of blanks and primers used can be found in Supplementary Table 1.

### Bioinformatic pipeline and taxonomic assignment

The raw sequencing data (13 615 653 606 reads) were processed using a customised version of the ‘HOLI’ pipeline [39] by Liu et al. [34]. In short, quality checked reads were deduplicated before and after trimming from adapters, followed by alignment against a custom database using Bowtie2 [40]. The taxonomic classification was performed using metaDMG [42], and ngsLCA [43]. The custom reference database was generated from RefSeq [44] and NT January 21 databases from NCBI [45], PhyloNorway contigs [45], Bacterial and Archaeal genomes of GTDB v220 [47], high-quality virus genomes of IMG/VR v4.1 [48], and selected genomes from selected taxa of the Siberian (sub)arctic.

### DNA damage pattern analysis and authentication

Post-mortem DNA damage was assessed using the PyDamage (v0.72) pipeline as described in [49]. Briefly, quality-filtered, error-corrected metagenomic reads were de novo assembled into contigs, aligned, and processed to estimate damage profiles. Contigs were taxonomically classified with Kraken2 [49] against the nt database (downloaded in October 2022), and damage estimates were merged by contig ID. We then applied PyDamage [50] to assess post mortem damage patterns for viruses and bacteria using contigs with a prediction accuracy ≥0.6 and length ≥1000 bp. 5′ C-to-T substitution frequencies were summarised by taxonomic group. Reads mapping to these retained contigs were considered as potentially ancient.

### Data analysis

All subsequent data analyses were performed using R (v4.3.2; https://www.r-project.org/) with tidyverse (v2.0.0) [51]. Unless otherwise stated, plots were generated using ggplot2 (v3.5.1) and cowplot (v1.1.3; https://wilkelab.org/cowplot/). The maps with the coring locations were generated using sf (v1.0-16) and rnaturalearth (v1.0.1). Plot alignments were done using Affinity Designer (v1.10.8).

To account for differences in sequencing depth, taxonomic count data were normalised by total sum scaling, yielding relative abundances. Rarefaction was not applied due to the low number of viral reads and the risk of losing rare taxa. The ngsLCA output is provided as read counts assigned to individual taxa, where reads attributed to higher taxonomic levels do not include those assigned to subordinate taxa. Hence, for each sample, reads assigned to lower-level taxa (species, genus, family, order, class, phylum) were cumulatively aggregated, yielding read counts at all taxonomic levels.

To explore the overlap of taxonomic diversity across sediment cores, virus species lists were extracted for each core and compared and visualised in an upset plot using ggVennDiagram (v1.5.3) [52]. In total, the reads assigned to viral species account for 94.7%.

Community structure was further assessed using non-metric multidimensional scaling (NMDS) based on Bray–Curtis dissimilarities of Hellinger-transformed relative abundances of the virus species. Redundancy analysis (RDA) and partial RDA were performed on Hellinger-transformed relative abundances using the vegan package (v2.6.4; https://vegandevs.github.io/vegan/) to evaluate the contribution of environmental variables and geological epochs on the variance of the community composition. Statistical significance of the constrained ordinations was assessed using permutation-based ANOVA as implemented in the *anova.cca* function from the vegan package and 1000 permutations.

Temporal patterns in virus community composition were visualised using stratigraphic area plots, with relative abundances with respect to viral reads plotted against calibrated ages. Only taxa with consistent presence or high relative abundance were included in stratigraphic summaries.

To assess long-term virus–host co-occurrence patterns, we performed correlation analyses based on cumulative relative abundances, calculated with respect to all reads. Viruses were associated with their host based on published literature and databases (Supplementary Table 2). Viral species were grouped according to their inferred host groups, and their relative abundances were summed accordingly. For host taxa, eukaryotic hosts were summarised at the phylum level, whereas bacterial hosts were summarised at the family level. This taxonomic resolution was chosen based on the assumption that eukaryotic viruses often exhibit broader host ranges than bacteriophages.

In a complementary analysis focused specifically on bacteriophage–bacterium dynamics, viruses were grouped according to the bacterial class of their known or inferred host. Host information was derived from a curated virus–host pairing list, supplemented by the Virus Metadata Resource by ICTV (VMR/MSL38 v2; https://ictv.global/news/vmr_release_0923).

When virus–host associations were not explicitly listed, we used the trivial virus names, which typically include the host genus followed by “phage”, to infer host identity. Viral genera were then assigned to host bacterial classes based on their known host genus.

### Data availability

The sedaDNA shotgun sequencing raw data is stored at the European Nucleotide Archive under the following Bioproject IDs: Lake Levinson Lessing (PRJEB82351), Lake Lama (PRJEB80877), Lake Ilirney (PRJEB80635), Lake Bolshoe Toko (PRJEB80642), Lake Ulu (PRJEB82635), PS97/72-01 (PRJEB74305), SO201-2-77KL (PRJE866300), SO201-2-12KL (PRJEB46821).

The documentation of the taxonomic assignment of sedaDNA shotgun data of the HOLI pipeline is available under https://github.com/sisiliu-research/EnviHoli. The analysis code is available at https://github.com/christianeboe/sedaDNAviruses (will be deposited on Zenodo upon acceptance).

## Results and discussion

### Ancient viruses in lake and marine environments

We retrieved a total over 13 billion (13 615 653 606) paired-end reads from 254 lake and marine sediment samples and 69 blanks across eight sites (Figure 1A). Of these, 400 186 289 reads (2.93%) aligned to eukaryotic, bacterial, archaeal, or viral DNA genome references of our custom database. Viruses accounted for 357 161 reads (0.089% of all mapped reads), across 2084 identified virus taxa and a median of 47 virus taxa and 396 reads per sample. Of all viral reads, 94.7% were assigned at species level, representing 65.3% of the virus taxa, whereas ∼4% could not be subclassified within the Virus realm. The remaining viral reads were assigned to genus level or higher.

**Figure 1:**
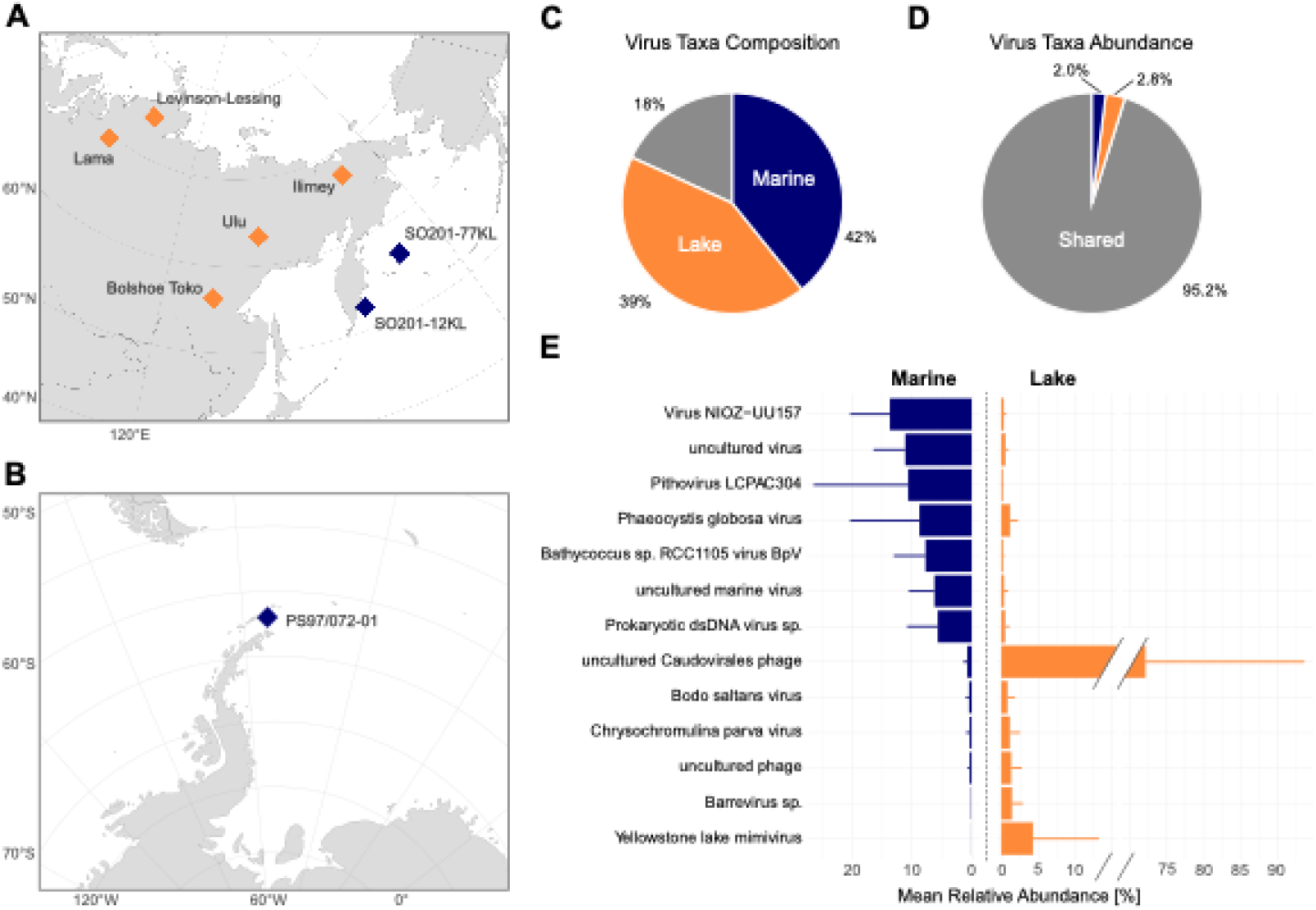
Site locations and virus species overlap. **A** Map of the coring sites in the **A** Northern and **B** Southern Hemisphere. **C** Proportion of viral species unique to marine or lake environments, or shared between both. **D** Proportion of reads assigned to taxa unique to marine or lake environments, or shared between both. **E** Bar plot of mean relative abundance (± standard deviation) of the most abundant shared taxa. Colours indicate lake (orange), marine (blue), and shared (dark grey) environments.

We then applied PyDamage [50] to assess post-mortem damage patterns for viruses and bacteria. We filtered contigs with a prediction accuracy ≥0.6 and length ≥1000 bp and plotted 5′ C-to-T substitution frequencies for each taxonomic group. Reads mapping to these retained contigs were considered as potentially ancient. These contigs exhibit characteristic ancient DNA damage patterns, particularly elevated C-to-T substitution rates at the 5’ ends. Average rates at the first nucleotide position range from 0.12–0.19 for viruses and are comparable to those seen in bacterial hosts (0.11–0.19). Among damaged reads, a pronounced decline in C-to-T frequency across the first 10 read positions is observed (Supplementary Figure 1 and 2), consistent with expected post-mortem damage accumulation. Notably, the C-to-T frequency is generally lower at site PS97, a pattern also observed in other taxa from this site [57], likely reflecting better overall DNA preservation in this sediment core. Taken together, the high number of ancient contig assignments and consistent damage patterns support the conclusion that both viral and host DNA are indeed ancient.

Most virus species are unique to either lake (879; 43%) or marine (745; 39%) sediment cores, with only 382 species (18%) shared between the two environments (Figure 1C).

These unique taxa are rare, comprising just 4.8% of viral reads, while the shared species account for 95.2% but are typically dominant in only one environment (Figures 1D, 1E). This

separation is also apparent in non-metric multidimensional scaling (NMDS), where samples cluster by environment (Figure 2A), consistent with modern aquatic viromes in which marine and freshwater systems share only a small fraction of virus taxa [10]. This pattern may reflect the rarity of marine–freshwater transitions for viruses and their hosts, due to osmotic barriers, competition, and predation by locally adapted species [50].

**Figure 2:**
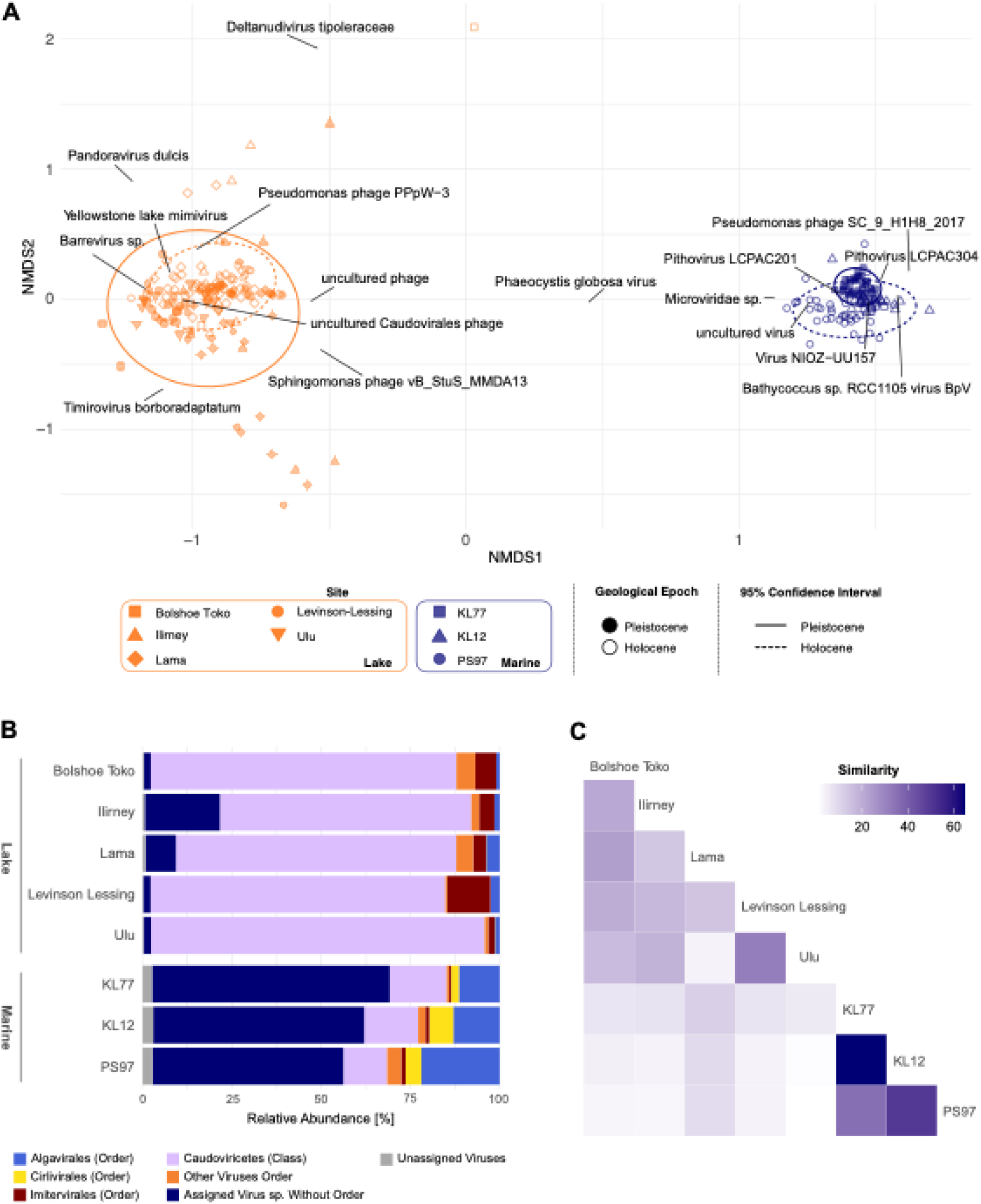
Virus community composition. **A** Non-metric multidimensional scaling (NMDS) plot of virus species composition coloured by environment: marine (blue) and lake (orange). Individual sites are distinguished by shape; Pleistocene samples are filled, and Holocene samples are empty. Ellipses represent the 95% confidence intervals for Holocene (dashed line) and Pleistocene (solid line) samples. Loadings of the 16 most abundant virus species are labelled. **B** Composition of ancient virus orders in lake and marine cores. The three most abundant virus orders are coloured light blue, yellow, and red. Caudoviricetes (aggregated at the class level) are shown in purple. All other orders are grouped and shown in orange. Viruses classified below the superkingdom level but unassigned to an order are in dark blue. Viruses not classified within the virus superkingdom are shown in grey. **C** Heatmap of pairwise similarity of the virus community between sites. Darker shading indicates higher similarity.

Samples are also grouped by geological epoch (Holocene vs. Pleistocene) in the NMDS ordination, based on ages determined through radiocarbon dating (Figure 2A). Similar differences between glacial and interglacial virus communities have been observed in ice core studies [32], although one permafrost study reported no strong differences, likely due to coarser taxonomic resolution [27]. In modern samples, polar and temperate viral communities differ substantially in both lakes [11] and oceans [8,12]. Hence, the observed epochal differences here are likely linked to the temperature gradient between the geological epochs.

Redundancy analysis (RDA) confirms that both environment and epoch significantly influence viral community composition, jointly explaining 50.1% of the variance (p < 0.001). Partial RDA shows that environment alone accounts for 49.0% (p < 0.001), while epoch explains 0.5% (p < 0.001), indicating a weaker but statistically significant effect. These patterns are consistent with environmental drivers in modern systems [8,10–12] and support the use of sedaDNA to reconstruct ancient viral assemblages.

Each environment is characterised by a distinct viral community. Lake samples are dominated by *Caudoviricetes*, which infect bacteria and archaea, and many viruses belong to *Imitervirales*, known to target amoebae (Figure 2B). In marine samples, a large proportion of viruses could be assigned at the species level but remain unclassified in the current ICTV taxonomy. Among classified groups, *Caudoviricetes* and *Megaviricetes* are most abundant. The latter includes *Algavirales* and *Imitervirales*, which target algae and protists, respectively, consistent with modern marine samples [1]. *Cirlivirales,* a group with a broad host range including chordates, crustacea, and possibly seaweed [51], are notably more abundant in marine samples (median = 3.41%) than in lake samples (median = 0.29%).

In 86% of lake samples (117 of 136), over half of viral reads are assigned to NCBI Taxonomy ID 2100421, labelled as ‘uncultured Caudovirales phage’. This group, comprising over 4000 environmental DNA sequences classified within the *Caudoviricetes* class, is predominantly associated with freshwater environments. It is highly dominant (>90% of viral reads) in 26% of lake samples but reaches only 4% in marine samples (Supplementary Figure 3). Given that only two sequences in this group originate from estuarine marine contexts, their presence in marine sediments likely reflects allochthonous input from terrestrial and freshwater sources, similar to patterns observed for plant sedaDNA [52].

Ancient marine viral communities are more similar across sites than those in lakes (Figure 2C). In particular, the two North Pacific cores KL12 and KL77 are highly similar, which coincides with a high number of shared virus taxa (Supplementary Figure 4). In contrast to the marine cores, which share 94 virus taxa, the lakes exhibit more distinct virus communities, with only 23 shared taxa (1.1%). Each lake harbours 66–226 unique virus taxa, more than are shared with other lakes (e.g. 42 shared between Lakes Lama and Levinson-Lessing). This mirrors high virus endemism observed in contemporary polar lakes, likely driven by limited connectivity [11], and parallels elevated host endemism in polar and high-elevation lakes [52]. The strong similarity between marine cores, particularly those in close proximity, reflects the connectivity of global ocean water masses and suggests viral dispersal via ocean currents, as has been demonstrated in modern viromes [8,10].

### Host-driven differences in ancient viral community structure across marine environments

Virus species composition differs considerably between the Antarctic core PS97/72-1 and the subarctic Pacific cores SO201-2-12KL and SO201-2-77KL (hereafter PS97, KL12, and KL77). In the NMDS analysis, samples from PS97 are spatially separated from KL12 and KL77 (Figure 2B). The distinct virus community in PS97 is driven by a high relative abundance of alga-infecting viruses, while KL77 and KL12 contain more viruses infecting bacteria. The abundance of alga-infecting viruses in PS97 coincides with a high relative abundance of algae [53], resembling contemporary Southern Ocean communities [54].

During the ACR, PS97 is dominated by *Pithovirus* LCPAC304 (mean relative abundance: 21.2%), which declines to 0.96% after the ACR. This species is unclassified, and its host is unknown. It is described as Pithovirus-like due to sequence similarity with *Pithoviridae* genes [55], suggesting it may infect *Pithoviridae* hosts such as amoebae or other protists.

After the ACR, the Haptophyta-infecting genus *Prymnesiovirus* becomes dominant (Supplementary Figure 5). This shift coincides with a transition from a Haptophyta- dominated to a *Chaetoceros*-dominated diatom community, based on analyses of the same DNA extract [53]. Sea ice and temperature have been identified as potential drivers.

Although the role of viral infection was not tested, *Prymnesiovirus* activity may have contributed to establishing a new post-ACR steady state.

Around 8 cal ka BP, *Prasinovirus* became abundant, coinciding with an increase in its host *Bathycoccus prasinos* and a shift towards warmer open-ocean conditions [56]. In modern samples, *B. prasinos* is associated with polar waters but is less abundant in temperate regions [57]. However, growth may have been restricted in the colder conditions that prevailed in the Bransfield Strait before 8 cal ka BP [56].

In contrast, *Prasinovirus* and *Prymnesiovirus* are abundant in KL77 and KL12 throughout the late Pleistocene but decline at around 12 and 14 cal ka BP, respectively (Supplementary Figures 6 and 7). Prior to the LGM, KL77 also has high abundances of *Pelagivirus* and *Stopavirus*, both of which infect *Pelagibacter*, a member of the ubiquitous SAR11 clade. With the onset of the Holocene (∼14 cal ka BP), *Kyanoviridae*, including *Llyrvirus* and *Mazuvirus*, both targeting *Synechococcus*, become abundant.

### Temporal shifts in lake virus communities since the Last Glacial Maximum

Temporal changes in lake viral communities are highly variable across sites. While many abundant taxa are present across multiple sites, their temporal patterns differ markedly, even in lakes that are geographically close or share climatic zones (Supplementary Figures 8–12). Despite site-specific differences and the persistent dominance of *Caudoviricetes*, all lakes exhibit temporal shifts from the late Pleistocene to the Holocene. For the last glacial time-span, *Casjensviridae* are detected in all lakes except Levinson-Lessing, probably being derived from the catchment, as their hosts are associated with terrestrial plants (Supplementary Figures 8–12). Although relatively rare (1.0–4.6%), they occur only before ∼10 cal ka BP, making them the only virus family more abundant in cold than warm periods. Their absence in Lake Levinson-Lessing may be due to colder conditions that suppressed host presence or viral transmission. The dominant genus, *Salvovirus*, infects *Xylella*, a widespread plant pathogen that relies on insect transmission and is positively associated with higher temperatures [59]. *Xylella* is present in all lakes during the glacial, indicating cold tolerance, but elevated *Casjensviridae* abundance during this time implies that infection may have been favoured under extreme conditions. The Holocene sees a rise in alga-infecting *Phycodnaviridae*, especially in Lakes Levinson-Lessing and Ilirney (Supplementary Figures 8–12), likely reflecting enhanced aquatic productivity [59,60].

*Mimiviridae* comprise up to 40% of viral reads in Lake Levinson-Lessing and up to 10% in the other lakes, making it one of the most abundant families. Its abundance remains stable or declines with Holocene warming in Bolshoe Toko, Lama, Ulu, and Ilirney. In contrast, it increases more than sixfold in Lake Levinson-Lessing after 13.5 cal ka BP, becoming dominant alongside *Caudoviricetes*. *Mimiviridae*, the main representative of *Imitervirales* in lakes (median = 100%), is slightly less abundant in polar than temperate lakes [11], which may indicate a temperature-dependent distribution. While this pattern could explain the Holocene increase, it contrasts with the higher abundance in Levinson-Lessing relative to warmer, lower-latitude lakes.

Interestingly, Lake Levinson-Lessing shares more viral taxa with marine cores than any other lake in the dataset (24 taxa; Supplementary Figure 4), which have higher abundance before 44.2 cal ka BP (Supplementary Figure 13), consistent with palaeogeological evidence of marine inundation of the Taymyr Peninsula during the Middle Weichselian [62]. Therefore, the virus community at the time likely reflected the more marine conditions that were present during the inundation period.

### Virus–host co-occurrence patterns on millennial timescales

On millennial timescales, virus and host abundances are mostly positively correlated in both lake and marine environments (Figure 3A). For this correlation analysis, we identified 83 virus–host pairs for the most abundant virus species, based on databases and peer-reviewed literature. These pairs involve 28 host taxa spanning all domains of life. We calculated correlations using the cumulative relative abundance of viruses infecting the same host. Correlation coefficients are significantly greater than zero in both marine and lake systems (Wilcoxon signed-rank test, one-sided, p < 0.001). This is in line with our expectation that hosts must co-occur with their viruses on long-term scales, because sediment samples integrate occurrences of several decades. As the sampling resolution reflects variation on multi-millennial timescale, short-term virus–host dynamics cannot be resolved.

**Figure 3:**
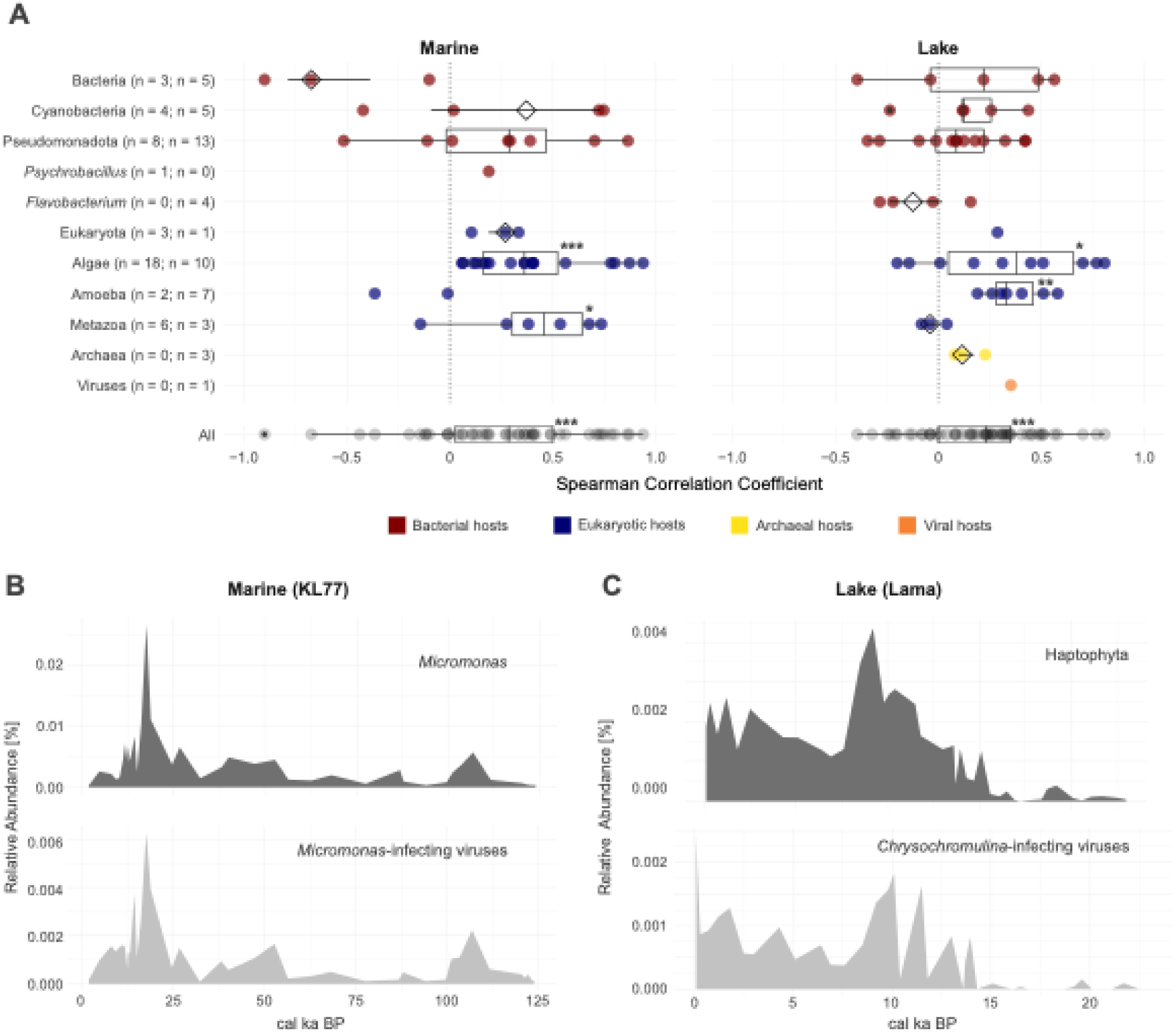
Abundance correlations of virus–host pairs. **A** Virus–host abundance correlations in marine (left) and lake (right) environments. Each point represents the Spearman correlation coefficient of a virus–host pair at a specific site. The number of correlations per environment is indicated. Host abundances were aggregated at the genus level for bacteria (red), the phylum level for eukaryotes (blue), and the superkingdom level for archaea (yellow). All correlations are summarised in a boxplot (black) at the bottom. **B** Temporal relative abundances of *Micromonas* (top) and *Micromonas*- associated viruses (bottom) at site KL77. **C** Temporal relative abundances of Haptophytes (top) and associated viruses (bottom) at Lake Lama. Relative abundances are shown as a proportion of total reads. Stars above boxplots indicate significance levels from one-sided Wilcoxon signed-rank tests (p ≤ 0.05 (*), p ≤ 0.01 (**), p ≤ 0.001 (***)).

Correlations are particularly strong between algal hosts and their viruses (Figure 3A). For example, *Micromonas* and its viruses follow the same temporal patterns across 125 cal ka in core KL77 (Figure 3B). In lakes such as Lama, we observe strong correlations between *Chrysochromulina*-infecting viruses and the broader Haptophyte phylum, which includes *Chrysochromulina* (Spearman = 0.81; Figure 3C). In aquatic systems, abundances of phytoplankton and their viruses are often tightly coupled, because they follow seasonal bursts and collapses during phytoplankton blooms [62]. Hence, strong correlations between them on longer timescales are expected.

### Introduction of Arctic-origin host strains and disrupted virus–host dynamics after the Bering Strait opening

Many abundant virus species from the North Pacific marine cores belong to *Pelagivirus*, which infects Pelagibacterales (SAR11), the most abundant bacterial clade in the oceans [63]. In KL77, *Pelagibacter*-infecting viruses are strongly correlated with their host in the Pleistocene (Spearman = 0.983) but not in the Holocene (Spearman = 0.103). This shift is driven by a single strain, *Candidatus* Pelagibacter sp. IMCC9063, which becomes highly abundant around 12.4 cal ka BP and sharply declines in abundance after 4.7 cal ka BP (Figure 4). This strain is not correlated with any identified *Pelagibacter*-infecting virus in KL77 (Supplementary Figure 14). Removing it restored the correlation between the remaining *Pelagibacter* species and their viruses, resulting in a strong positive Spearman coefficient of 0.855 (p = 0.003) also in the Holocene samples.

**Figure 4:**
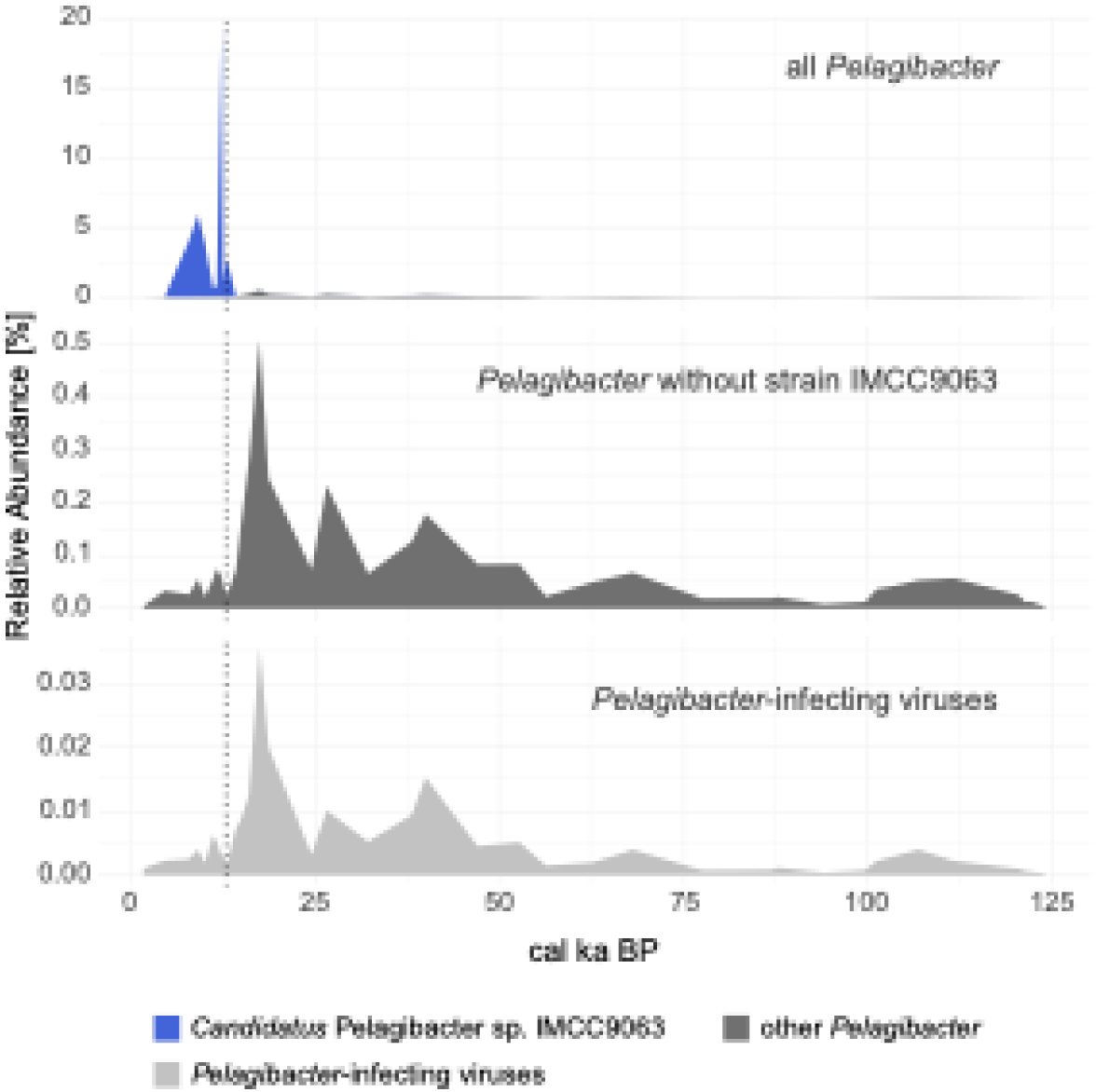
Abundance patterns of *Pelagibacter* and associated viruses. Temporal relative abundances of *Pelagibacter* and *Pelagibacter*-infecting viruses at site KL77. The upper plot shows all *Pelagibacter* species and strains, with strain IMCC9063 highlighted in blue and all others aggregated in dark grey. Due to the high relative abundance of IMCC9063, other *Pelagibacter* species are nearly undetectable in this plot. The middle plot displays *Pelagibacter* species excluding IMCC9063 (dark grey), revealing temporal dynamics of less dominant taxa. The lower plot shows *Pelagibacter*-infecting viruses (light grey). The dotted line indicates the timing of the Bering Strait opening.

A similar pattern is seen for *Synechococcus* (Supplementary Figure 15). The strain *Synechococcus* sp. Ace-Pa dominates the genus around 12.4 cal ka BP and decreases in abundance shortly after. It skews the correlation analysis suggesting a lack of correlation between virus and host (Spearman = -0.176, p = 0.785), but when excluded, the virus–host correlation becomes positive for the Holocene (Spearman = 0.527, p = 0.003).

The abundant *Pelagibacter* and *Synechococcus* strains are both associated with polar waters. *Candidatus* Pelagibacter sp. IMCC9063, isolated from the Arctic Ocean [64], is limited to waters <18.2°C [69] and carries cold-adaptation genes under selection in Arctic and Antarctic SAR11 strains [69]. *Synechococcus* sp. Ace-Pa, isolated from a marine Antarctic Lake [70], has not yet been found elsewhere, though a related strain may also occur in the Arctic.

Despite favourable glacial conditions, both strains become abundant only after the Bering Strait opened (∼11–12 ka BP) [18,19]. This suggests their introduction from the Arctic Ocean into the Bering Sea following the first inundations of Beringia. No identified *Pelagibacter*- infecting virus mirrors the host peak at this time, indicating the absence of viral regulation. Although the absence of a respective virus species in this dataset does not imply a general lack of viral infection of these strains at the time, their strong dominance in the bacterial community supports the hypothesis that extensive viral lysis did not occur. Their decrease in relative abundance after 9 cal ka BP may be explained by environmental changes such as increasing sea surface temperatures [71,72] which became less suitable for cold-adapted hosts.

### Mismatch between bacteriophages and bacterial host class composition

The most abundant bacteriophages do not match the dominant bacterial classes in marine or lake systems (Figures 5A and 5B). Pleistocene marine communities are dominated by Gammaproteobacteria (Figure 5A), in line with contemporary marine environments [73], but their viruses are rare. Actinobacteria are the only group to co-occur with abundant viruses in Holocene marine cores KL77 and KL12. In comparison, Betaproteobacteria and Alphaproteobacteria occur in lakes (Figure 5B), consistent with modern observations [74,75].

**Figure 5:**
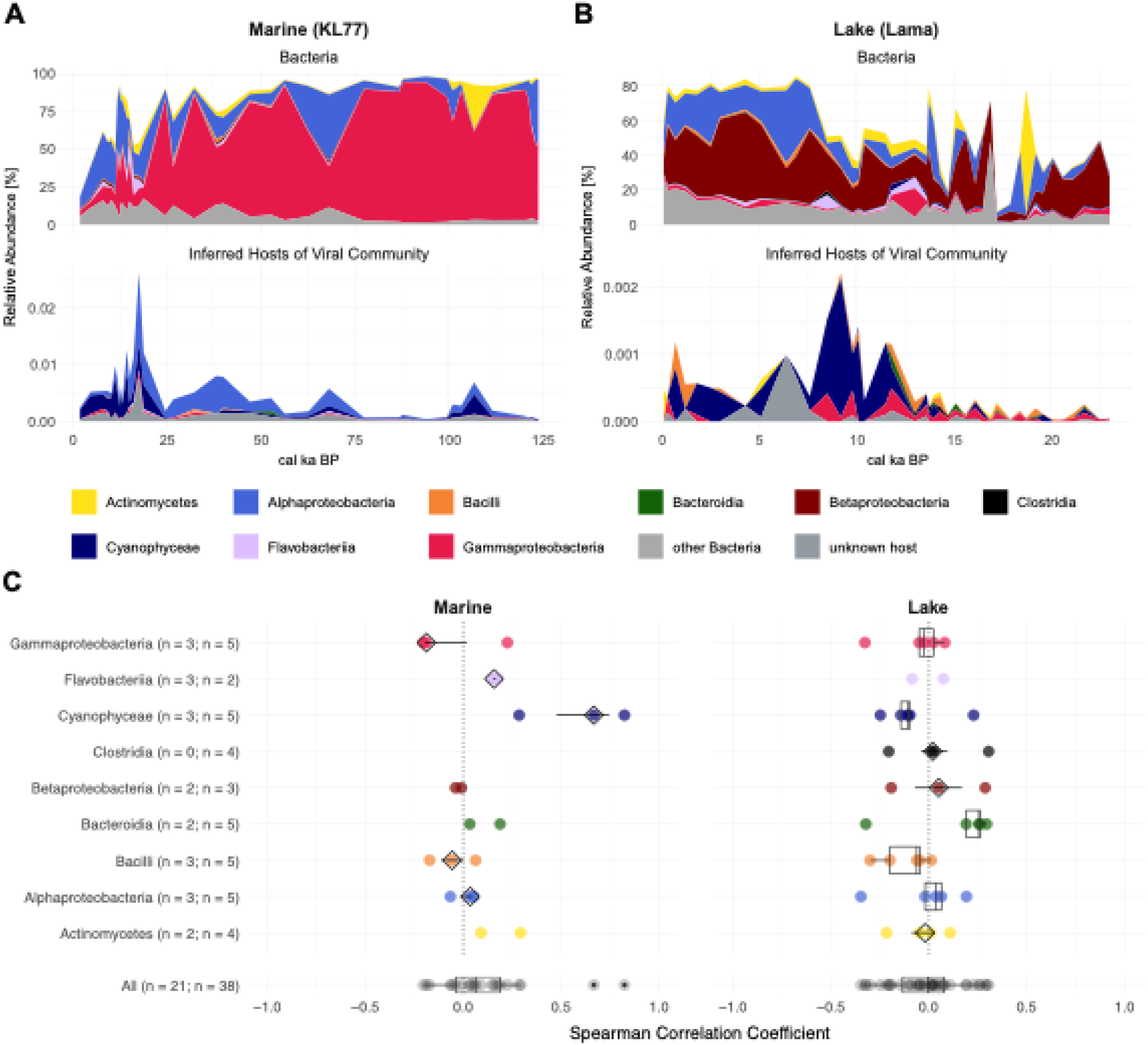
Ancient bacteria and their associated bacteriophages. **A T**emporal relative abundances of bacterial classes (top) and their associated bacteriophages (bottom) in cores KL77 and **B** Lake Lama. The colours indicate the bacterial classes and the inferred hosts of the associated viruses. **C** Boxplots of Spearman correlation coefficients between bacterial classes and associated bacteriophages in marine (left) and lake (right) environments. Each point represents the correlation at a specific site. Samples are coloured according to bacterial class (as in A and B). Combined distributions of all correlations are shown in black at the bottom of each panel.

However, phage communities in both systems are dominated by cyanobacteria-infecting viruses, while their hosts, Cyanophyceae, are rare (marine: 0.10%, lake: 0.76%) (Figures 5A and 5B).

Despite the low relative abundance of Cyanophyceae, they correlate strongly with their viruses in the marine system but not in the lakes (Figure 5C). All other highly abundant bacterial classes and their viruses do not correlate in either lake or marine environments. In lakes Lama, Bolshoe Toko, and Ulu, Cyanophyceae and their viruses even show antagonistic patterns (Supplementary Figures 16–18), reflected by negative correlation coefficients (Figure 5C). These antagonistic patterns can also be observed between the whole bacterial and bacteriophage communities at Lake Ilirney (Supplementary Figure 19) as well as in marine cores KL12 (Supplementary Figure 20) and KL77 (Figure 6A).

Additional negative correlations include *Pseudomonas* in Lake Ilirney (Supplementary Figure 21) and Gammaproteobacteria in lakes Bolshoe Toko and Levinson-Lessing and subarctic marine cores KL12 and KL77 (Supplementary Figures 22–25).

While short-term antagonism fits the Kill-the-Winner model [66], long-term data should show covariation due to cumulative signals. Instead, these antagonistic patterns on millennial timescales may result from other factors. It is important to note here that during periods of high virus abundance, their host abundance is not zero, indicating that the host is present but at low relative abundances. Ongoing viral infection and lysis of a host group allows organisms to succeed in the competition for growth resources, which shifts the composition in favour of the organisms that experience less pressure from viruses [76]. Therefore, an increase in abundance of a virus can coincide with host declines due to competition.

Conversely, reduced viral pressure can allow host populations to rebound. These shifts may reflect co-evolutionary dynamics described by the Red Queen hypothesis [9]. However, antagonism may also reflect viral shifts between lysogenic and lytic cycles [4,77,78].

Lysogenic viruses produce little viral DNA [79], making them harder to detect in sedaDNA datasets. Integrated viruses may even benefit hosts relative to competitors [76]. Yet, when the viruses revert to a lytic lifestyle, the host can no longer establish dominance within their community and they decline in relative abundance. This phenomenon of suppressed host dominance through viral lysis may explain both antagonistic temporal patterns and mismatches between dominant bacteriophages and bacteria.

The only exception to this mismatch are Actinobacteria in marine environments, which comprise the highly abundant SAR11 clade, including the *Pelagibacter* genus. The success and ubiquitous abundance of the SAR11 clade was previously attributed to viral resistance [2]. However, this theory was later disproved due to the high abundance of Pelagiphages [67]. The high relative percentage of Pelagiphages throughout the late Quaternary implies a sustained virus production, presumably reliant on host lysis, and thus challenges the viral- resistance hypothesis.

These findings illustrate that virus–host correlations are not consistently positive and that decoupled or antagonistic patterns may reflect ecologically meaningful processes. It is therefore important to consider that negative correlations between viruses and their hosts do not necessarily indicate false associations, but may reflect shifts in infection dynamics, particularly across environmental transitions such as glacial-interglacial warming. Such dynamics may be more common over long timescales than previously assumed and should be accounted for when using co-occurrence-based approaches to infer host-virus relationships in ancient datasets.

## Conclusions

This study is the first to demonstrate the feasibility of using sedaDNA to reconstruct ancient DNA virus communities in both lake and marine environments, providing new insights into long-term virus–host dynamics across geological epochs. Our results show that temporal patterns in ancient viromes are preserved and biologically meaningful, despite low proportions of viral reads.

By comparing lake and marine viromes, we find that distinct viral community signatures persist over millennial timescales and through major environmental changes. This persistent separation underscores the importance of ecosystem-specific processes in shaping viral assemblages. While marine viral communities are more interconnected, likely due to ocean circulation, lake communities are more isolated and show high levels of endemism, including a marine-like viral signature in Lake Levinson-Lessing, likely reflecting inundation during the Middle Weichselian. These findings align with contemporary patterns and extend them back through glacial-interglacial transitions.

Viral community composition differs between glacial and interglacial periods, with environment (lake vs marine) explaining most of the variance, followed by temperature. However, additional factors such as long-term evolutionary dynamics may also influence viral assemblages, as shown by the distinct virus composition during MIS5, which resembles the subsequent glacial more than the Holocene. Lakes also have greater inter-site dissimilarity than marine systems, supporting the idea that local conditions play a dominant role in shaping freshwater viral communities.

Temporal structure reveals clear ecological responses to climatic shifts. Lakes show shifts in dominant viral taxa at the onset of the Holocene, including increases in algal viruses with rising aquatic productivity, and the disappearance of cold-associated phage families like Casjensviridae. In marine systems, transitions such as the Antarctic Cold Reversal and the postglacial opening of the Bering Strait coincide with marked shifts in virus–host associations. These patterns suggest that ancient viral communities tracked both local and global environmental changes across time.

Using a literature-based approach, we identified 83 virus–host pairs and find that most exhibit positive correlations in their relative abundances, consistent with long-term co-occurrence. These correlations are particularly strong for phytoplankton and their viruses, reflecting tight ecological coupling. However, we also detect non-correlated and antagonistic patterns, some of which we could link to geological events (e.g., Bering Strait opening) or hypothesised life-cycle switches between lytic and lysogenic phases in response to environmental change. Our results highlight the importance of considering negative correlations between viruses and their hosts when applying co-occurrence-based approaches to infer virus–host relationships. Such antagonistic patterns may be common over long timescales, especially across warm–cold transitions or other major environmental changes that affect infection dynamics.

Despite these promising insights, several challenges remain. Interpretation of ancient viromes remains limited by a high proportion of unassigned reads, restricted viral reference databases, and the current focus on DNA viruses. Future studies should expand reference collections, pursue genome-resolved reconstructions, and aim to integrate RNA viruses.

This study shows that sedaDNA offers a powerful tool to investigate ancient virus–host ecology, revealing how these interactions evolve over long timescales and respond to environmental change. As modern ecosystems face accelerating climate shifts, these long-term perspectives offer a valuable baseline for understanding microbial community dynamics in changing environments.

## Acknowledgements

We acknowledge Cathy Jenks for manuscript proofreading. AI assisted copy editing has been applied using ChatGPT (https://chat.openai.com/).

